# A draft atlas of the human plasma proteome at isoform resolution

**DOI:** 10.64898/2026.09.26.754256

**Authors:** Boris Martin Zühlke, Torsten Schwecke, Tal Weizmann Shapira, Agathe Niewienda, Anyi Yang, Fatma Amari, Kathrin Textoris-Taube, Ludwig Roman Sinn, Karola Lehmann, Christoph Gille, Martin Vingron, Michael Mülleder, Markus Ralser

## Abstract

Key plasma protein, including albumin, complement system proteins, and coagulation factors circulate in biochemically and functionally distinct isoforms, or proteoforms. Despite the importance of these proteoforms being widely accepted, they are challenging to resolve systematically, and as a consequence, we still lack a more complete picture of the total proteoform diversity of the human plasma proteome. Here, we generated 3,114 intact protein fractions by extensively separating plasma according to isoelectric point and molecular weight. We then analysed these fractions by liquid chromatography data-independent acquisition mass spectrometry (DIA-MS) upon tryptic digestions, and combined the resultant data into a multidimensional, proteoform resolved atlas of the human plasma proteome. We report that 5,077 unequivocally detected plasma proteins circulate at least 179,049 distinct proteoforms, of which we detect 44,083, with a median of 11 proteoforms per protein, at high confidence. Substantially increasing the number of experimentally confirmed proteoforms, our atlas implies that the canonical protein forms account for less than half of the total protein mass in plasma. We detect the greatest proteoform diversity among the most abundant plasma proteins and those functioning in nutrient transport, complement and coagulation. Finally, our data clarifies discrepancies between the Olink and SomaScan proteomic platforms, as according to our atlas, these techniques provide correlated values for relatively abundant proteins with low to medium proteoform diversity, but struggle in the quantification of low-abundant proteins and on those which exist in many proteoforms. In sum, our data reveals extensive proteoform diversity of the human plasma proteome, and implies that proteoform diversity represents an enormous untapped reservoir for biomedical research and biomarker discovery.

## Introduction

The blood plasma is a human tissue uniquely specialized in transport, signal transduction, and immune mediation, and as such a central component of physiology. The plasma connects tissues and organs through the transport of nutrients, metabolites, hormones and lipids, while simultaneously supporting immune defense, coagulation, and systemic homeostasis (Schaller et al. 2008; Kennelly et al. 2015; Hall and Hall 2025). Its accessibility and systemic nature have also made plasma and serum predominant biological matrices for clinical diagnostics. Many of the most widely used biomarkers in medicine are measured in serum or plasma, including protein markers, such as cardiac troponins for the diagnosis of myocardial injury, cystatin C for assessment of kidney function, and C-reactive protein as indicators of inflammation (Geyer et al. 2016; Carrasco-Zanini et al. 2024). Nevertheless, sensitive and specific biomarkers or biomarker panels remain unavailable for many diseases, particularly at early stages when therapeutic intervention may be most effective. Identifying new protein markers for early disease detection, patient stratification, treatment selection and monitoring therefore remains a major objective of biomedical research and an important prerequisite for precision medicine (Deutsch et al. 2021; Messner et al. 2023).

Thus far, the search for new biomarker strategies has often focussed on rendering lowly abundant proteins better detectable. In particular, highly multiplexed affinity-based platforms such as Olink proximity extension assays and SomaScan aptamer-based assays now enable large numbers of protein targets to be measured across tens of thousands of samples (Sun et al. 2023; Dhindsa et al. 2023; Eldjarn et al. 2023). These technologies have enabled population-scale studies linking circulating proteins to genetic variation, disease risk and clinical outcomes (Geyer et al. 2024; Sun et al. 2024; Carrasco-Zanini et al. 2024). However, comparisons between platforms have also revealed a fundamental limitation: measurements assigned to the same protein frequently show only modest or even poor agreement between technologies or replicates, despite the presumed intrinsically technical reproducibility (i.e., low variance) of the individual assays (Rooney et al. 2025; Pietzner et al. 2026). There are two main hypotheses to explain these difficulties. Firstly, physical limits apply to quantifying proteins over nine or more orders of dynamic range, irrespective of which methodology is used. It is thus plausible that such low correlation emerges from a limited quantification precision of proteins challenging to quantify. Secondly, since plasma proteins can exist in many proteoforms, these could be recognized differently by the affinity reagents. Indeed, all well-studied plasma proteins such as albumin, apolipoproteins or complement proteins occur as multiple proteoforms, generated through proteolytic processing, post-translational modifications (PTMs), truncation, alternative splicing, and other biochemical transformations (Tagliabracci et al. 2015; Mastellos et al. 2024; Chevalier et al. 2025). Albumin for instance, circulates in numerous modified and truncated forms; apolipoproteins undergo extensive proteolytic and post-translational processing and associate with distinct lipoprotein particles; and complement proteins are converted through regulated proteolytic cleavage into molecular species with distinct biological activities (Tagliabracci et al. 2015; Rhode et al. 2019; Demir et al. 2025). Consequently, assays assigned to the same gene or canonical protein sequence could in practice measure different subsets of the circulating protein population. Despite mass spectrometry detecting multiple peptides per protein, also in this field the dominating practice is to estimate a single protein value per protein (Cox et al. 2014). While this practice certainly works well for cellular and microbial proteomes and reduces noise, it results in an underutilisation of proteomic data in proteoform rich matrices, especially the human plasma proteome.

Due to these technical limitations, we still lack a map that captures a baseline, or ground truth, of the plasma proteome. Here, we address this gap by generating a draft version of a proteoform-resolved map of the human plasma proteome. We achieve this by excessively exploiting the power of classic biochemical protein separation techniques, isoelectric focussing (IEF), denaturing size exclusion chromatography (dSEC), and 2-dimensional polyacrylamide gel electrophoresis (2D-PAGE) to separate proteoforms by mass and charge (Andrews 1964; Dale and Latner 1969; O’Farrell 1975; Anderson and Anderson 1977). We combine this ‘top-down’ separation with the sensitivity of peptide-centric, ‘bottom-up’ proteomics, recording proteomes using liquid chromatography data-independent acquisition mass spectrometry upon tryptic digestion. Using this approach, we resolve approximately 5,000 human plasma proteins into 180,000 distinct proteoforms, revealing a previously inaccessible layer of molecular complexity within the human proteome for biomedical research and biomarker discovery.

## Results

### Identification of proteoforms in the human plasma proteome through the systematic separation of intact protein fractions

In order to generate intact protein fractions of a human plasma sample, we decreased the concentration of 14 most abundant proteins from a plasma sample by applying an affinity resin (Tu et al. 2010) (see Methods) and then fractionated the resulting intact protein fraction by i) iso-electric focussing, ii) denaturing size exclusion chromatography and iii) four replicates of two-dimensional polyacrylamide gel electrophoresis (2D-PAGE, see Figure 1, Methods). For IEF and dSEC we generated 24 and 18 samples, respectively, for proteomic analysis. For 2D-PAGE, every replicate gel was cut into 768 cubes of 8 x 8 x 1 mm (voxels). In order to analyse these by bottom-up proteomics, we automated the HiT-Gel approach for in-gel digestion of proteins (Swart et al. 2018). Furthermore, to every sample we added equal amounts of the Open Standard for Plasma Proteomics (OSPP), an internal, isotopically labelled peptide standard mixture of 211 consistently detectable heavy isotope labelled peptides that match major plasma proteins, herein used for normalisation and quality control (Wang et al. 2025). We analysed equal volumes of the gel-based peptide digests on an Evosep One liquid chromatography system (Evosep) coupled to a timsTOF HT mass spectrometer (Bruker) operated in diaPASEF acquisition mode (Meier et al. 2020). The resulting raw data files were processed with diaTracer (K. Li et al. 2025) within the FragPipe computational suite (Kong et al. 2017; Demichev et al. 2021) to generate an extensive plasma proteome spectral library containing tryptic and semi-tryptic peptides derived from the curated UniProt Proteome (UniProt Consortium 2025) and the PeptideAtlas database (Desiere et al. 2006) (Weizmann Shapira et al., parallel manuscript). Next, we analysed the mass spectrometric raw data with DIA-NN 2.3.1 (Demichev et al. 2020) using this extended spectral library for peptide precursor quantification. Precursors were filtered on precursor Q-values, global Q-values, library Q-values below 1% and PEP-values below 50% and summarised to peptide quantities employing the MaxLFQ algorithm (Cox et al. 2014; Pham et al. 2020). After scaling every peptide across every fractionated dataset to values between 0 and 1 we removed 3% of samples marked as outliers based on the presence of contaminations and OSPP quality assessment (see Methods). This pre-processing created a digital representation of peptide quantities for IEF, dSEC and the replicate 2D-PAGE experiments. In order to identify proteoforms, within these 1D and 2D projections, we identified peaks for every peptide above a peptide-specific noise threshold, defined as the median peptide abundance plus two times the standard deviation of its noise level (Immerkær 1996) and 10% above neighbouring sample quantities using the peak_local_max algorithm from the python scikit-image package (van der Walt et al. 2014). The set of peptide peaks within a protein group was considered to represent the proteoform landscape of each protein group. To match proteoforms per protein group across 2D-PAGE replicates, we determined node-assignment integer linear programming problems on distance matrices reduced to matchings within replicate-specific thresholds, decomposed into connected components solved independently with the CP-SAT solver from the python OR-tools package version 9.12 (Google 2026).

**Fig. 1:**
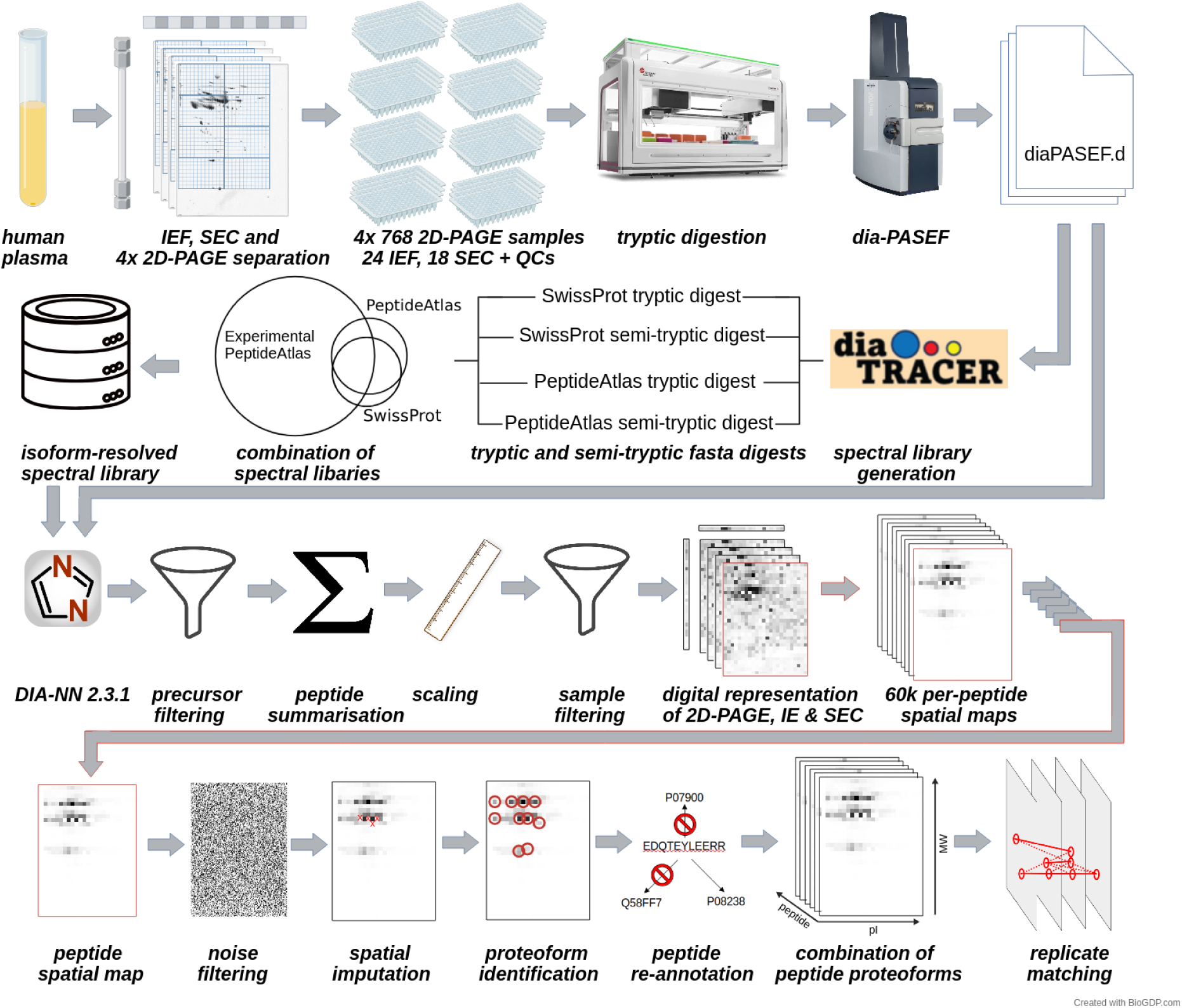
Mapping the proteoform diversity of human plasma by intact protein separation followed by bottom-up proteomics. A human plasma sample was separated by denaturing size exclusion chromatography (dSEC), iso-electric focusing (IEF), and in four replicate 2D-PAGE experiments. The intact protein fractions were then analysed by bottom-up mass spectrometry, using diaPASEF, DIA-NN, and a spectral library generated by combining spectral-centric database searches on the obtained raw data with information from the PeptideAtlas resource (Kong et al. 2017; Demichev et al. 2020; K. Li et al. 2025). A data analysis pipeline recreated the digital representations of the fractionated datasets and per-peptide proteoforms were identified and annotated to their respective canonical protein groups. Confidence for protein identification was established at the peptide precursor level, and confidence of proteoform differentiation by comparing the digital representation of the separation experiments. Proteoforms detected in at least three out of four 2D-PAGE experiments were considered high-confidence proteoforms. Created with BioGDP.com (Jiang et al. 2025).

Our strategy distinguished 177,894 proteoforms for 4,687 protein groups detected across four 2D-PAGE experiments. Of these we identified 44,083 proteoforms for 2,089 detected protein groups in at least three out of the four replicate 2D-PAGE experiments. Herein we refer to this set as the high-confidence proteoform map. One-dimensional separation by dSEC and IEF, respectively, resulted in 7,700 proteoform detections for 2,492 protein groups detected by IEF and 6,343 proteoforms for 2,150 protein groups detected by dSEC. Most of these were also detected in the 2D-PAGE dataset (Figure 2A). Notably, the better separation using 2D-PAGE, which identified twelve times more separable proteoforms than both 1D-dimensional separation techniques together (Figure 2A and 2B), is largely explained by the increased separation power of the 2D-PAGE experiment: projecting the identified proteoforms from the 2D-PAGE onto one dimension also resulted in 7-fold more proteoforms compared to IEF and 10-fold more compared to dSEC (Figure 2B). Summarizing all datasets, we detected in total 179,049 proteoforms for 5,077 protein groups containing 4,440 unique proteins aggregating to 3,671 proteins with unique protein matching. For a group of 314 predominantly highly abundant proteins, we confirm 4,083 proteoforms at maximum confidence: these 4,083 proteoforms were matched between all four replicate 2D-PAGE experiments, dSEC and IEF runs, and detected by at least three peptides in every 2D-PAGE replicate experiment.

**Fig. 2:**
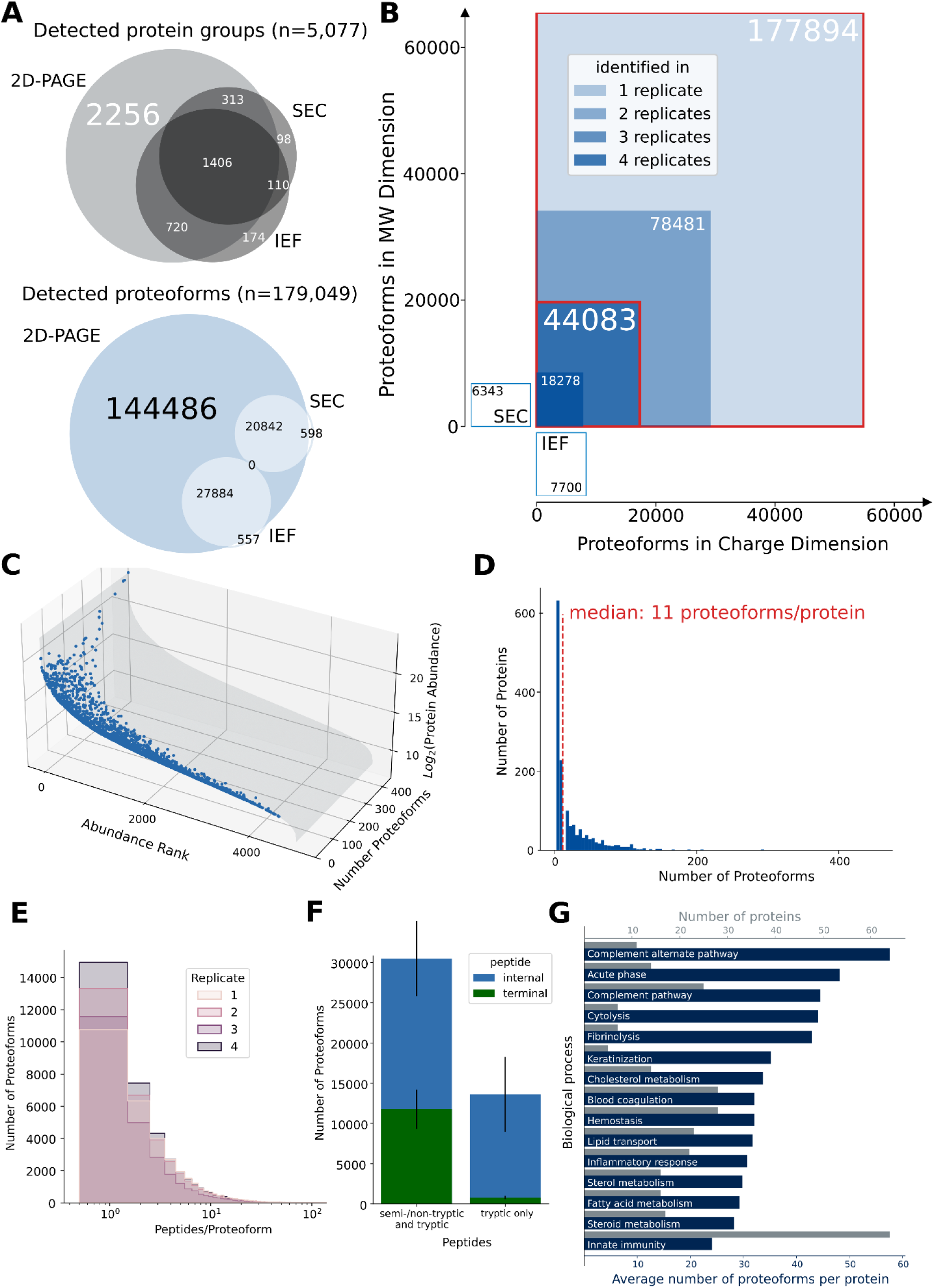
The human plasma proteome is characterized by a massive proteoform diversity. A: Identified protein groups (upper Venn diagram) and identified proteoforms (lower Venn diagram) using 2D-PAGE, denaturing size exclusion chromatography (dSEC) and iso-electric focusing (IEF) followed by diaPASEF; 2D-PAGE experiments were projected to their charge and size dimension to be compared to dSEC and IEF, separately, thus a proteoform from the 2D-PAGE separation may be matched to both, dSEC and IEF with a possible shift of max. 4 fractions as obtained from image overlay of the replicate gels; the total numbers count unique protein groups/proteoforms. B: Numbers of identified proteoforms scaled per dimension with annotated confidence as number of gels a proteoform was identified in. The higher the confidence, the darker the colour. The set of any identified proteoform (complete proteoform map) and proteoforms identified in at least 3 out of 4 experiments (high-confidence proteoform map) are marked with red boxes. C: Protein groups ranked by their abundance in the 2D-PAGE data with the number of identified proteoforms annotated in the third dimension from the high-confidence proteoform map (Scatterplot). D: Distribution of proteoforms per protein group for the high-confidence proteoform map (Histogram). E: Distribution of proteoforms by their number peptides they are identified with across the four replicate 2D-PAGE gels for the high-confidence proteoform map. F: Number of proteoforms with tryptic, semi-tryptic and non-tryptic peptides versus proteoforms identified by only tryptic peptides across the high-confidence proteoform map for protein sequence-internal (blue) and -terminal (green) peptides; error bars indicate the standard deviation of proteoform numbers across replicate 2D-PAGE gels. G: Enrichment analysis on the top 15 Human Protein Atlas - Biological Process terms with highest number of identified proteoforms/protein in high-confidence proteoform map excluding potential contaminants compared to the number of proteins for the respective terms (Álvez et al. 2025).

On the other hand, numerous proteoforms were identified by only one peptide, many of which are lowly abundant proteins (Figure 2C and 2E). Nonetheless, after matching proteoforms between the replicate 2D-PAGE experiments, only 11% of high-confidence proteoforms were detected by just one peptide across replicates; in other words, 89% of repeatedly detected proteoforms were identified by multiple peptides, and thus likely represent true positive identifications.

### Characterizing the proteoform landscape of the plasma proteome

High-confidence proteoforms were observed across the detection range (Figure 2C), and on average, our dataset lists eleven high-confidence proteoforms per protein group (Figure 2D). For the maximum confidence set (mostly highly abundant proteoforms, detected in six out of six experiments with a minimum of three peptides detected in every experiment for every proteoform) we call 4 proteoforms per protein. We noted that most proteoforms were detected among the highly abundant proteins. This has in part technical reasons, as lowly abundant peptides are more difficult to detect, but it seems not to be an overall artefact, as also abundance of proteins in the mid-abundance range correlates with their number of proteoforms, and is consistent with the notion that many highly abundant proteins function in processes for which many proteoforms are already known, such as metabolite transport, coagulation and complement activity. To gain more insight of the protein abundance on the number of proteoforms, we calculated knee and elbow points using the protein abundance ranks to classify proteins into highly abundant (n=663), mid-range (n=3,981) and lowly abundant (n=212) (Figure 4F). For the mid-range proteins, we observed a median of five proteoforms per protein, while for the highly abundant proteins this increased to 29 proteoforms per protein. For 405 protein groups detected in neat plasma (Wang et al. 2025), we identified nine proteoforms per protein group for mid-range abundant proteins, and for the top 44 proteins classified as highly abundant, this number increases to 50 proteoforms. To identify the biological processes related to most of our detected proteoforms, we then counted the occurrence of *Biological Process* (BP) terms in the Human Protein Atlas annotation (Uhlen et al. 2010) linked to protein groups and proteoforms (Figure 2G). Here, the most frequent BP terms found were complement alternate pathway, acute phase and complement pathway. In contrast, for the BP term innate immunity, we found lower proteoform diversity (Figure 2G). Thus, despite acknowledging technical limitations in the detection of lowly abundant protein isoforms, our data suggests that highly abundant plasma proteins contribute most to proteoform diversity in human plasma.

As there is currently no gold standard for the presence of proteoforms in human plasma, it is difficult to estimate how many new proteoforms are discovered by our dataset. However, we compared our high-confidence proteoform map to the aggregated UniProt database, wherein proteoform diversity can be extrapolated from isoform annotations, variants and post-translational modifications (UniProt Consortium 2025; Kalyuzhnyy et al. 2022). For 71% (1,384 proteins), our map identified more proteoforms than protein isoforms listed in UniProt for the particular protein. Furthermore, despite the presence of a PTM site or a sequence polymorphism not necessarily constituting a distinct proteoform (e.g. many PTMs, such as phosphorylations, show interdependency, and many proteins detected in cells are not excreted into the plasma), we identify more proteoforms for 72% and 66% of proteins than number of reported PTMs and variants, respectively. Along these lines, we compared our number of identified proteoforms to the sum of isoforms, PTMs and variants reported for the same proteins on UniProt. Also compared to this number, our map identifies nearly 150% more proteoforms as present in UniProt (Figure 3C).

**Fig. 3:**
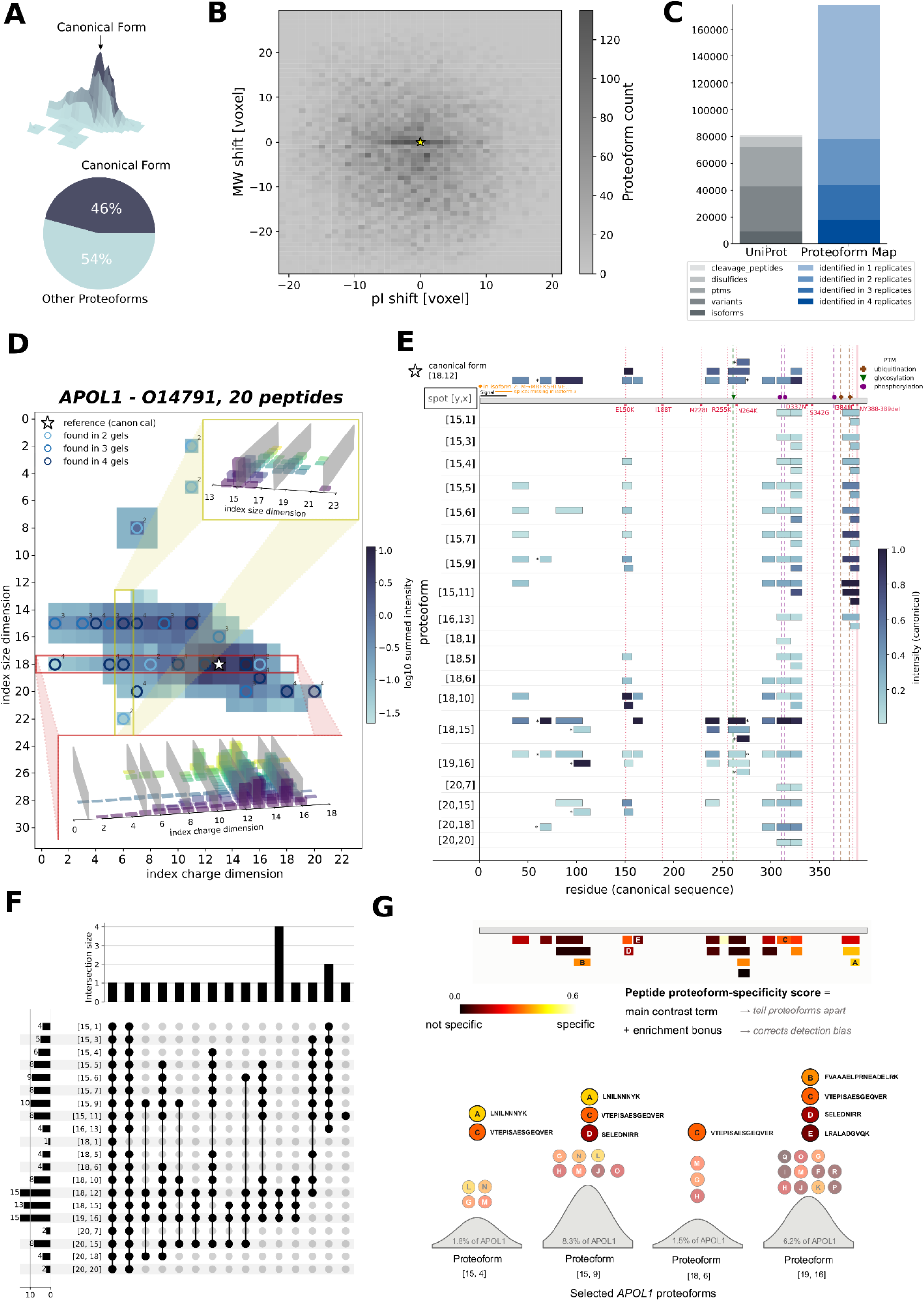
Plasma proteoforms are characterized by specific peptide patterns. A: Total signal of a protein group subdivides into the most abundant proteoform (here defined as the canonical form) and all other proteoforms. B: Distribution of proteoform shifts relative to the canonical form (centered at [0,0], marked by the star) within one replicate 2D-PAGE example. C: Comparison of the number of proteoforms identified across 2D-PAGE replicas in the high-confidence proteoform map to the UniProt database concerning each proteins’ annotated number of isoforms, variants (natural SNPs), ptms (post-translational modifications), disulfides and cleavage peptides (Ahmad et al. 2025). D: 2D-PAGE representation for the protein APOL1 in an exemplary 2D-PAGE experiment with identified peaks; darker colour represents higher scaled peptide intensity sum, detected proteoforms are marked with circles and the most abundant proteoform (canonical protein) is marked by a star; circle colour and annotated number represents the number of matched replicate gels. Per peptide abundance profiles across size and pI dimension are shown as insets on a selected dimension in red and yellow, respectively. Proteoform peaks in these 3D barplots are represented as grey rectangles across peptides. E: Peptide coverage across proteoforms (rows with index coordinates in D) for the protein APOL1 on one replicate 2D-PAGE gel. Peptides are represented by rectangles at their respective position in the amino acid sequence, darker colour represents higher peptide abundance and asterisks mark non-tryptic cleavage sites; on top the canonical proteoform and sequence with known modifications (glycosylation = green, phosphorylation = purple, ubiquitination = brown), natural variants (red) and splice locations (orange) collected from the UniProt database is shown. F: Upsetplot demonstrating the peptide overlaps across the 20 proteoforms detected in the high-confidence proteoform map for APOL1. G: Peptide coverage and proteoform-specificity scoring for APOL1. Top panel: overall peptide coverage of the canonical APOL1 sequence, in total 50% of the sequence are covered by detected peptides; peptides are coloured by their proteoform-specificity score, with yellow indicating greater and black indicating lower discriminatory power between proteoforms. Lower panel: proteoform specificity score using four selected APOL1 proteoforms ([15, 4], [15, 9], [18, 6], [19, 16]) and five representative peptides (A: LNILNNNYK, B: FVAAAELPRNEADELRK, C: VTEPISAESGEQVER, D: SELEDNIRR, E: LRALADGVQK) spanning the score range.

### Assessing the source of proteoform diversity

Next, we performed an initial assessment of whether a proteoform originated through sequence modifications such as proteolytic cleavage or alternative splicing, or through post-translational modifications of the canonical sequence. One indicator for sequence-altered proteoforms is the presence of non-tryptic cleavage sites originating from biological processing. Our map assigned peptides with non-tryptic cleavage sites to two thirds of the high-confidence proteoforms (Figure 2F, blue). Additionally, proteoforms identified with non- or semi-tryptically cleaved peptides are overrepresented by terminal peptides (Figure 2F, green).

To characterize proteoforms relative to their canonical protein identity we then defined the most abundant proteoform by summed scaled peptide intensities as the canonical form (H.-D. Li et al. 2014). Our data indicates that less than 50% of the summed measured peptide quantity of a protein group arises from its canonical proteoform and more than half of the measured intensity is distributed to less abundant proteoforms in our map (Figure 3A). We also detect distinct groups of proteoforms shifted to the canonical form along the isoelectric point dimension with no effects on the mass dimension (most likely PTMs), as well as many smaller proteoforms for most proteins, most likely originating from alternative splicing and proteolytic cleavage (Figure 3B).

As an example to demonstrate the resolution of our proteoform map we turned to APOL1, an apolipoprotein from the apolipoprotein L family (UniProt Consortium 2025). APOL1 has a predicted protein mass of 44 kDa, assembled from 398 amino acids, and features a predicted iso-electric point (pI) of 5.5 (Kozlowski 2022). The UniProt database reports for APOL1 three isoforms (including the canonical form), seven single nucleotide genetic variants (SNPs), one glycosylation site, two ubiquitination sites and three phosphorylation sites. Considering the distribution of plasma proteins by abundance, APOL1 is a mid-abundant protein, ranked 490^th^ by abundance in our proteoform map and 145^th^ out of 405 proteins in an LC-MS based proteomic dataset generated from neat plasma (Wang et al. 2025).

Our map suggests that APOL1 is present in many more proteoforms as a projection from the coding sequence may propose. APOL1 was identified by multiple peptides, at least 14 and a total of 25 peptides, across the individual separation experiments. We illustrate summed scaled peptide intensity of APOL1 within one replicate 2D-PAGE experiment (Figure 3D). APOL1 was detected over a broad abundance range, spanning 9 out of 32 voxels on the size dimension (dimension ranges from 150-10kDa) and 22 out of 24 in the pI dimension (dimension ranges from pI 3-10) (Figure 3 D). Across all experiments we detected 41 APOL1 proteoforms, of which 20 were identified across at least 3 out of 4 replicate 2D-PAGE separations (27 in the illustrated example, Figure 3D). APOL1 proteoforms can be grouped into two major size variants (Figure 3D). We highlight the proteoforms by circles colour-coded with the confidence assigned through the number of replicates in which the proteoform and the dominant (canonical) form (marked with a star, Figure 3D) were detected. The peptide composition across proteoforms along one dimension is illustrated in an inlaid 3-dimensional barchart showing individual peptide abundances across the given dimension. Although not all proteoforms can be distinguished by summarised protein abundance alone, peptide profiles across the 2D-PAGE experiment provide distinct proteoform-discriminating peptide footprints (Figure 3D-F).

Mapping identified peptides onto the amino acid sequence of the canonical form (Figure 3E grey, top) shows an altered peptide composition across APOL1 proteoforms (row positions in Figure 3E) which imply different length variants (first index of a tuple) that underlie some of the proteoforms. For example, proteoforms from the smaller major size variant (size index 18 and higher) all lack the peptides starting at positions 372, 373 and 381, indicating that this region is missing in the small proteforms, while the peptide starting at position 373 uniquely distinguishes proteoform [15,11] (Figure 3E and 3F). Other proteoforms [18,15] and [19,16] which have a similar iso-electric point are distinguished by peptides starting at position 255, which carry a conserved glycosylation site on N261 (Liu et al. 2005). For these two peptides, we can also identify the natural variant N264K that introduces a tryptic cleavage site at position 264. Moreover, our APOL1 proteoform map identified proteoforms with non-tryptic cleavage sites which could mark potential alternative splicing events or proteolytic cleavage sites so far unreported in literature (Figure 3E). Thus, the example of APOL1 illustrates a proteoform diversity of a typical, mid-abundant plasma protein and demonstrates the assignment of specific peptide patterns to the 2D-PAGE separated isoforms. In our parallel manuscript by Weizmann Shapira et al., we use this information to annotate proteoform specificity to individual peptides using a deep spectral library (Weizmann Shapira et al., parallel manuscript) (Figure 3G). Exemplified by APOL1 we show how our proteoform specificity scoring incorporates relative differential abundance of peptides across proteoforms by a contrast term and corrects for generally low abundant peptides by an additional enrichment term (Figure 3G). Our peptide proteoform specificity score successfully assigned higher proteoform specificity to peptide A, which exhibits a strong differential abundance pattern across proteoforms of higher size and is absent among proteins of lower size (Figure 3E and 3G). Peptide B, which is specific to a small group of spatially close proteoforms and more abundant in proteoform [19,16] than in the generally more abundant proteoform [18,15], also has a higher proteoform specificity score assigned. In contrast, the peptide E which is only found in one highly abundant proteoform but generally lowly abundant itself got correctly assigned a low proteoform specificity. Peptides D and C, which are omni-present across many proteoforms, possess less proteoform discriminatory power and consequently have lower proteoform-specificity scores assigned (Figure 3E and 3G).

### The role of proteoform diversity for the quantification of the plasma proteome with affinity reagents

It is a common concern that several proteomic methods currently applied to human plasma in cohorts disagree in their reported protein quantities, even when applied to the same sample set. This situation puts uncertainty in genetic linkage studies such as genome wide associations, the use of artificial intelligence which is very sensitive to data quality, as well as biomarker discovery programs. For example, in the Atherosclerosis Risk in Communities Study (THE ARIC INVESTIGATORS 1989; Rooney et al. 2025), plasma proteomes of 102 study participants were measured in duplicates on the SomaScan 11k (v5.0, SomaLogic) and Olink Explore HT (Olink Proteomics) platforms. The technologies reported quantitative values on 9,658 and 5,417 unique UniProt Ids, respectively; however, the quantities obtained were found to be in poor overall agreements, both in the cross-platform comparison, but also in duplicate measurements using the same technology (Rooney et al. 2025). Only 2,100 abundance values quantifying 1,202 proteins (UniProt Ids) were at least in moderate (R > 0.4) agreement between both technologies as well as the respective replicates (Figure 4A and 4C). Only a minority of 144 protein quantities for 73 proteins (i.e., UniProt Ids) agreed well (R > 0.9). Furthermore, from the 10,784 proteins measured by SomaScan 11k, just 59% agreed well (R > 0.8), and 37% agreed very well (R > 0.9) in duplicate measurements. Similarly, for the 5,420 Olink-derived Ids measured by Olink Explore HT only 37% agreed well between duplicate measurements (R > 0.8) and merely 25% agreed very well (R > 0.9) (Figure 4A and 4D).

**Fig. 4:**
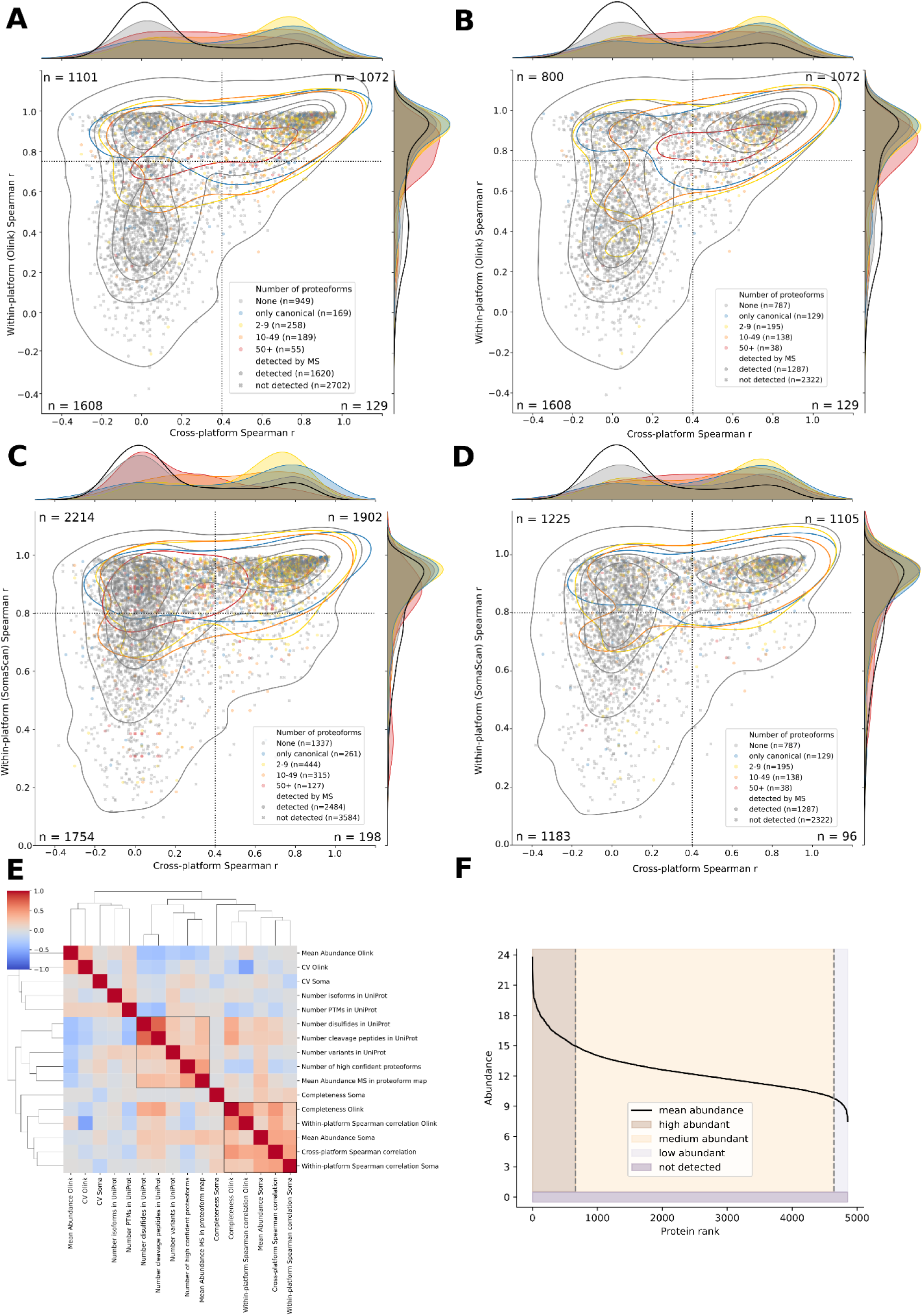
Proteoform and protein abundance and their relationship with quantification precision in Olink Explore HT and SomaScan 11k assays. A-D: Dependence of Olink Explore HT (A,B) and SomaScan 11k (C,D) within-platform correlation (between duplicate measurements of patient samples) to cross-platform correlation between protein abundances measured on the SomaScan 11k and Olink HT Explore platform for 6,068 protein Ids in the Atherosclerosis Risk in Communities Study (ARCS) across 102 participants (Rooney et al. 2025); A and C show all measured proteins per UniProt Id, B and D only the protein with maximum correlation within a UniProt Id; scatter style marks if proteins were detected in our experiments by bottom-up LC-MS/MS, marker colour defines the number of proteoforms identified for the detected proteins, n-values in corners count the number within every quadrant obtained by the separation by correlation thresholds 0.4 for cross-platform correlation and 0.75 and 0.8 for within-platform correlations (Olink Explore HT and SomaScan, respectively). Marginal histograms (kernel-density estimates) represent the distribution of proteins (black = not detected by LC-MS/MS, grey = detected by LC-MS/MS and coloured according to the number of identified proteoforms (Rooney et al. 2025). E: Spearman correlation clustermap of summary statistics in the ARCS, proteoform information from the UniProt database and number of proteoforms and protein abundance within our high-confidence proteoform map. F: Classification of protein abundances into high, medium and low abundance by elbow and knee criteria. The mean abundance is based on LC-MS-based proteomics measurements.

Here, we used our proteoform map to shed more light on which protein quantities agree or disagree between platforms and duplicates (Figure 4). Plotting the abundance values generated visually distinct protein clusters; protein quantities which agreed reasonably well between technologies and replicates (top right corner), and those which had poorer or no correlation. Firstly, proteins which were detected not only by Olink and SomaScan but also by mass spectrometry in our proteoform map, had a higher chance to agree between both affinity technologies and the Somascan replicates (p-value = 1.9 × 10^-78^, point biserial Pearson correlation = 0.24). Secondly, this group is significantly overrepresented by proteins with a low number of proteoforms (Fisher exact test for overrepresentation on protein Id bins with 1, 2-9 and 10-49 proteoforms, respectively, with Bonferroni multiple testing correction). Next, we focussed on the group of proteins that were in disagreement between the affinity reagent measurements. A smaller fraction of these proteins were similarly abundant as the consistently quantified proteins, but present in many (i.e., 50+) isoforms (Figure 4A-D). A larger fraction, however, was not detected at all by mass spectrometry (n=2,520), despite our extensive pre-fractionation, or detected at low abundance. In contrast, a third group of 519 proteins did agree between the affinity reagent measurements in the Atherosclerosis Risk in Communities Study and were not detected by mass spectrometry in our 2D-fractionated plasma sample. These proteins include disease-induced proteins not present in our baseline plasma sample. Thus, at least in this study, the main factor for disagreement between Olink and Somalogic measurements, as well as the reason for disagreement between replicates within the same technology, seems to be poor precision in the quantification of lowly abundant proteins, while in some cases, the disagreements is explained by reasonably abundant proteins that occur in many proteoforms. Considering only the highest cross-platform correlation per UniProt Id, we observe similar distributions regarding proteoform diversity, except that proteins with more proteoforms tend to be more uniformly distributed across the cross-platform correlation axis, and thus not overrepresented among proteins with low agreement between SomaScan and Olink (Figure 4B and 4D).

## Discussion

In this study we attempted to resolve the proteoform space of the human plasma proteome. Our approach was motivated by the success of classic biochemical separation techniques, SEC, IEF and 2D-PAGE in separating different molecular forms of the same, intact protein (Andrews 1964; Dale and Latner 1969; Anderson and Anderson 1977). Indeed, in the classic literature, 2D-PAGE provided some of the clearest visual evidence for proteoform heterogeneity, with individual proteins appearing as queues or constellations of spots rather than as single molecular species (O’Farrell 1975; Anderson and Anderson 1977). However, the limited sensitivity and throughput prevented this principle from being extended to the (plasma) proteome at scale. Here, we revisited this classical biochemical concept by expanding it with bottom-up proteomics technology using DIA-MS, applying a combination of HitGel (Swart et al. 2018), EvoSep One separation (Bache et al. 2018) and diaPASEF acquisitions (Meier et al. 2020), as well as advanced processing of the generated dataset with DIA-NN (Demichev et al. 2020), and a spectral library that combines the PeptideAtlas resource with additional spectra mapped with diaTracer and Frapipe (K. Li et al. 2025; Kong et al. 2017). In this way, we combine the resolving power of intact-protein fractionations of classic biochemistry, with the sensitivity, throughput and identification depth of contemporary bottom-up proteomics. Since 2D-PAGE is very effective at separating proteoforms, we conducted one of the most comprehensive protein fractionations experiments of the current literature, in which a reference sample was separated at the intact protein level, and analysed in 3,300 LC-MS/MS experiments.

We report a massive molecular diversity at the plasma proteoform level for 5,000 detected protein groups. Considering only the high-confidence proteoforms detected in at least three out of four independent separate experiments, our data reveals that the average plasma protein exists in at least 11 distinct isoforms (median), totalling to 40,000 high-confidence and almost 200,000 total proteoforms. Comparing the number of proteoforms identified in our analysis to reported numbers of isoforms, variants or post-translational modifications in the UniProt database suggests that we detected substantially more proteoforms than previously annotated.

Indeed, the unique biological function of the blood plasma renders protein diversification into proteoforms a plausible evolutionary solution leading to higher efficiency. While within cells, proteins predominantly have structural, catalytic, regulatory and organizational functions, in plasma, by contrast, protein function is dominated by molecular transport, intercellular communication, and circulation-specific functions like coagulation and complement activity (Jacobs et al. 2005). Especially transport activity depends strongly on molecular interactions driven by binding affinities. Proteoform diversity provides this functional variability without requiring the selection for new protein domains or new genes. For example, proteolysis or splicing can remove, add or alter interaction domains so that the similar protein scaffold can reach different tissues or bind different molecules for transport. Also proteolytic processing can expose or eliminate binding- or targeting sites. Post-translational modifications can further reversibly modulate molecular recognition, stability and interactions with receptors, ligands or carrier molecules. Truncations and other processing events may alter circulating half-life, clearance pathways or tissue targeting. It follows that for capturing the biology of the plasma proteome, it does not suffice to sum the proteforms into a single value that corresponds to the gene or origin or the shared sequence. Instead, technology is required to separate them into individual proteoforms.

Reporting an architecture of the human plasma proteome as a proteoform universe has important technical consequences for biomedical studies and biomarker discovery. Thus far, attempts to expand plasma proteomics for better biomarker discovery and biomedical studies have often focussed on the detection and quantification of lowly abundant proteins. While improving the detection of lowly abundant proteins certainly remains a powerful strategy, the proteoform diversity quantified herein has received far less attention. Indeed, our data suggests that there might be more distinct molecular species just among the highly abundant plasma proteins, as there are genes in the genome. Furthermore, less than half of the total plasma proteomic mass is contained within the most dominant form of the average protein. The potentially vast proteoform space thus constitutes a huge future opportunity for plasma biomarker discovery.

Using the mid-abundant protein APOL1 as an example, we demonstrate how our proteoform atlas can distinguish variations on the amino acid sequence level as well as on the post-translational modification landscape. We detect known natural variants and identify proteoforms previously not reported in literature. Furthermore, our first draft of a proteoform atlas provides scores for peptides, indicating how effectively they can distinguish proteoforms. This paves the way for differential analysis at the proteoform level in bulk proteomics studies of cohorts, eliminating the need for sample prefractionation.

Despite extensive fractionation and deep mass-spectrometric analysis in 3,300 LC-MS/MS experiments, we detected only approximately 5,000 protein groups. This number may feel comparatively small, given the massive analytical depth of the experiment and that higher numbers of proteins are detected by affinity methods. There are technical limitations to note, but also there are biological properties that dictate why many human proteins are not secreted at high concentration into the plasma, at least in a healthy individual. Technically, we have filtered our data to only include high-confidence peptides, and the separation of the plasma sample into thousands of fractions unavoidably dilutes the sample and may cause loss of some molecular species due to their physical properties. Indeed, despite our dSEC and IEF experiments creating far fewer fractions, they identified at least some proteins not detected in the 2D-PAGE fractions (Figure 2A). More importantly, however, seem to be the biological reasons that led also previous studies to consider similar numbers for the plasma proteome (Blume et al. 2020; Distler et al. 2025). First, the number of biological activities needed in plasma for human physiology is finite. Indeed, many enzymatic activities are undesired to occur in plasma. Enzyme activity can create reactive metabolites, interfere with immune cells or other tissues. Moreover, each protein leaked to plasma increases the risk of autoimmune reactions (Y. Li et al. 2025). Many proteins are thus prevented from leaking into the plasma at relevant concentrations for biological reasons. In this context, the comparison of our plasma proteome atlas with results obtained with the affinity reagent based Olink and Somalogic assays in the Atherosclerosis Risk in Communities Study (Rooney et al. 2025) yielded interesting results. Both Olink and Somalogic assays report values for many proteins that are not detected in our experiment. Indeed, specifically many of these proteins did not correlate between these two assay and replicate measurements (Rooney et al. 2025). This indicates that even if detected, these proteins are too lowly concentrated to be reliably quantified with these methods.

### Limitations and future directions

Several limitations of the present study define important directions for future work. Firstly, while our approach resolves a large number of distinct proteoforms according to their physicochemical properties, for most of these forms we do not yet know the molecular events responsible for their generation. Differences in apparent mass and charge arise through multiple mechanisms, including alternative splicing, proteolytic processing, post-translational modifications, or combinations thereof. The present map therefore establishes the existence and distribution of distinct molecular forms but does not, in most cases, provide their molecular characterization. Resolving this layer will require integration with complementary information, including peptide-level evidence for alternative sequences and cleavage events, modification-specific analyses, and orthogonal biochemical measurements. Particularly challenging forms will likely require sensitive top-down mass spectrometry to establish their intact molecular composition directly (Schaffer et al. 2019; Korchak et al. 2026; Rogers et al. 2026). Continued improvements in top-down proteomics, together with the substantial reduction in sample complexity provided by our fractionation strategy, as well as bottom-up proteomics data analysis approaches which leverage information on fragment ion level (Ammar et al. 2025), should make molecular annotation of an increasing fraction of the proteoform space feasible (Smith et al. 2021; Hollas et al. 2022; Bludau et al. 2021).

Secondly, our map remains constrained by analytical sensitivity. Although extensive prefractionation substantially increases the number of molecular forms that can be distinguished, detection ultimately depends on obtaining sufficient peptide-level signal from each proteoform. The current map is therefore expected to be most complete for abundant and intermediate-abundance plasma proteins, while the proteoform diversity of low-abundant proteins remains substantially underrepresented. This limitation may be particularly relevant for disease biology. Molecular forms that are absent or present at very low concentrations under physiological conditions may increase substantially during disease, tissue injury, inflammation or other perturbations. Such disease-associated forms would not necessarily be represented in a reference map generated from plasma of healthy individuals. Consequently, the 175,000 forms resolved here should not be interpreted as a fixed or exhaustive inventory of the human plasma proteoform space, but rather as a first, or draft, map of a molecular landscape that is likely to expand with increasing analytical sensitivity and across different physiological and pathological states.

Thirdly, the resolution of our fractionation systematically spans 8mm x 8mm squares of the majority of the 2D-PAGE gels. Proteoforms located close to the edge of a gel as well as larger proteoforms and signalling peptides were not included in the sample set. Image analysis reveals that many visible spots are split between samples or samples contain several spots which cannot be further distinguished. Some reported proteoforms are actually proteoform families understood as a set of similar proteoforms rather than distinct molecular forms. The effects of resolution in proteoform families remain to be studied.

Fourthly, our proteoform map includes all measured human proteins. The sample workflow itself is susceptible to contamination by cellular proteins (Geyer et al. 2019). Since many cellular proteins are assumed to be present in the plasma due to secretion or leakage from cells and tissues, and since known contaminant databases contain highly abundant plasma proteins such as albumin and complement factors, we decided to not exclude but flag potential contaminant proteins. Although the matching across replicate 2D-PAGE gels reduces the number of false positive proteoforms for contaminant proteins, our map does not provide confident false discovery rates on the proteoform level. As an additional result, our dataset extends contaminant lists with potential contaminants that could be identified across the majority of voxels of a 2D-PAGE experiment.

These limitations also highlight the opportunities created by a proteoform-resolved view of the plasma proteome. The present study thus provides a reference framework onto which molecular identities, inter-individual variation and disease associations can progressively be mapped. Combining such maps with increasingly sensitive top-down and bottom-up mass spectrometry, population-scale studies and clinical cohorts should ultimately establish which components of the extensive molecular diversity observed here represent constitutive plasma biology and which provide specific readouts of disease. In this sense, the current map represents a starting point rather than an endpoint towards understanding the human plasma proteome at the level of its functional molecular forms.

## Materials and Methods

### Immunodepletion of the plasma sample

A plasma sample was treated with the High Select Top 14 Abundant Protein Depletion Camel Antibody Resin (Thermo Fisher Scientific), in order to reduce the concentration of 14 most concentrated proteins. Prior to use, the resin was equilibrated from 4 °C to room temperature for 60 min, vortexed thoroughly, and aliquoted (300 µL per well) into 96-well deep-well plates (DeepWell™ Storage Plates, Waters). The plasma sample was diluted 1:10 (v/v) in 100 mM ammonium bicarbonate (NH₄CO₃), and 100 µL of diluted plasma were added to each well containing depletion resin. Plates were sealed with aluminum foil and incubated on a shaker (Heidolph Titrama× 101) at 1000 rpm for 1 h to ensure efficient mixing and antigen binding. Following a 10-min settling period, the supernatant was transferred to a fresh plate. The resin was subsequently washed with 500 µL of 100 mM NH₄CO₃, and the wash fraction was combined with the initial supernatant. Combined fractions were transferred to 96-well filter plates (Nunc™ 96-Well Filter Plates), and the plasma preparation was recovered by centrifugation at 100 × g for 2 min. into fresh deep-well plates. Samples were lyophilized and stored at −80 °C. Each well contained approximately 50 µg of protein. For 2D-PAGE, lyophilized proteins from ten wells (∼500 µg total protein) were reconstituted in 100 mM ammonium bicarbonate and desalted using 10 kDa molecular-weight-cutoff centrifugal filters (Merck). Protein concentrations were determined by bicinchoninic acid (BCA) assay (Thermo Fisher Scientific). Four independent 2D-PAGE experiments were performed by Proteome Factory (Berlin, Germany) on separate days.

### Isoelectric focusing (IEF)

Protein separation according to isoelectric point (pI) was performed using an OFFGEL Fractionator (Agilent Technologies) equipped with the OFFGEL Kit pH 3–10 and a 24-well setup (Agilent Technologies), following the manufacturer’s instructions. Immobilized pH-gradient (IPG) gel strips with a linear pH range of 3–10 were rehydrated in the assembled device using 40 μL of rehydration solution per well.

The plasma protein preparation containing 200 μg of protein was diluted in OFFGEL stock solution to a final volume of 3.6 mL. Subsequently, 150 μL of the diluted sample was applied to each well. IEF was performed at a maximum current of 50 μA and voltages ranging from 500 to 4,000 V until a total focusing voltage of 64 kV·h was reached. Focusing was completed after approximately 56 h. The recovered fractions, with volumes ranging from 100 to 150 μL, were subsequently processed using filter-aided sample preparation (FASP).

### Denaturing size-exclusion chromatography (dSEC)

Denaturing size-exclusion chromatography was performed using an ACQUITY UPLC H-Class system (Waters, Manchester, UK) equipped with a binary solvent manager, sample manager, column oven, and diode-array detector. Absorbance was monitored at 260 and 280 nm. Separation was carried out using a Yarra SEC-2000 column (300 × 7.8 mm, 3 μm particle size, 300 Å pore size; Phenomenex) at room temperature. The mobile phase consisted of 8 M urea, 1 mM dithiothreitol (DTT), and 10 mM sodium phosphate buffer adjusted to pH 7.0. Isocratic elution was performed at a flow rate of 0.5 mL/min. Protein samples containing 1 mg of total protein were injected, and fractions of 200 μL were collected.

### Filter-aided sample preparation (FASP)

Fractions of both, IEF and dSEC separations, were processed by filter-aided sample preparation (FASP) (Wiśniewski et al. 2009) using 30 kDa cutoff centrifugal filters in 96 well filter plates (Pall Corporation). Proteins were transferred to the filter units, and extensively washed with a urea-based buffer to remove contaminants. Following an additional wash with 50 mM ammonium bicarbonate buffer proteins were digested on the filter with sequencing-grade trypsin in a wet chamber at 37 °C overnight. The resulting peptides were collected by centrifugation and subjected to LC–MS/MS analysis.

### Sample preparation for two-dimensional polyacrylamide electrophoresis

During thawing of the samples, urea, carrier ampholytes, and dithiothreitol (DTT) were added to final concentrations of 9 M, 2%, and 70 mM, respectively. Samples were incubated for 30 min. at room temperature and subsequently centrifuged at 15,000 × g for 45 min. The resulting supernatant was transferred to fresh tubes and stored at −80 °C until analysis.

### Two-dimensional polyacrylamide gel electrophoresis (2D-PAGE)

Two-dimensional gel electrophoresis was carried by loading 90 µg of protein were loaded onto vertical rod gels containing 9 M urea, 4% acrylamide, 0.3% piperazine diacrylamide (PDA), 5% glycerol, 0.06% TEMED, 2% carrier ampholytes (pH 2–11), and 0.02% ammonium persulfate (APS). Isoelectric focusing (IEF) was performed to a total of 8,820 Vh.

Following IEF, gels were equilibrated for 10 min in buffer containing 125 mM trisphosphate (pH 6.8), 40% glycerol, 65 mM DTT, and 3% sodium dodecyl sulfate (SDS), and subsequently stored at −80 °C. For second-dimension separation, equilibrated IEF gels were applied directly to SDS-PAGE gels (20 × 30 × 0.1 cm) composed of 375 mM Tris-HCl (pH 8.8), 12.5% acrylamide, 0.2% bisacrylamide, 0.1% SDS, and 0.03% TEMED. Electrophoresis was performed at 140 mA for 5 h until the dye front reached the bottom of the gel. Proteins were visualized by silver staining (Proteome Factory, PS-2001).

### In-gel digestion and peptide extraction

Silver-stained gels were processed according to the HiT-Gel workflow described by Swart et al. with modifications (Swart et al. 2018). PAGE regions spanning a pI range of approximately 3–10 and a molecular weight range of approximately 15–150 kDa, encompassing all visible protein spots, were excised and subdivided into 768 sections (8 × 8 mm). Gel pieces were transferred into 96-well filter plates with temporarily sealed outlets and processed on a Biomek i7 (Beckmann) liquid handling workstation. Reagents were dispensed automatically, and liquids were removed by centrifugation following outlet release.

Proteins were digested overnight with Trypsin/LysC (Promega; 1 µg per sample). Peptides were extracted using 150 µL of 80% acetonitrile containing 0.1% formic acid, dried in a vacuum concentrator, and reconstituted in 40 µL of 0.1% trifluoroacetic acid (TFA). Peptide concentrations were determined using a fluorometric peptide assay (Pierce). Samples were spiked with 100 pg OSPP standard for sample quality assessment (Wang et al. 2025), and equal sample volumes were subjected to LC–MS/MS analysis.

### DIA LC–MS/MS analysis for one-dimensional separation techniques

Peptide analysis was performed using an Evosep One liquid chromatography system coupled to a timsTOF Pro mass spectrometer (Bruker). A total of 80 µL of peptide digest were loaded onto Evotip Pure tips according to the manufacturer’s instructions. Chromatographic separation was performed using the Evosep 30 samples-per-day method, employing a 44-min gradient on an EV1137 analytical column (15 cm × 150 µm, ReproSil-Pur C18, 1.5 µm particles; Dr. Maisch) maintained at 50 °C. The column was connected to a 10 µm Zero Dead Volume captive-spray emitter. Spectra were acquired across an *m/z* range of 100–1700 and an ion mobility range of 1/K₀ = 0.6–1.6 using 32 isolation windows per cycle. Collision energies were ramped according to ion mobility and ranged from 20 to 59 eV. The isolation window width was set to 50 Da, while accumulation and ramp times were maintained at 100 ms.

### DIA LC–MS/MS analysis for two-dimensional separation techniques

LC–MS analysis was performed using an Evosep One system coupled to timsTOF Pro (Bruker) mass spectrometer. Dried peptide extracts from the 2D-PAGE were resuspended in 80 µL 0.1% formic acid (FA) containing 100 pg OSPP peptides and loaded onto Evotip Pure tips according to the manufacturer’s instructions (Wang et al. 2025).

Liquid chromatography was performed using the Evosep 60 samples per day (SPD) method with a 21 min gradient on a PepSep column (8 cm × 150 µm, 1.5 µm particle size) maintained at 30 °C and coupled to a 20 µm zero-dead-volume captive spray emitter.

MS data were acquired over an *m/z* range of 100–1700 using an accumulation and ramp time of 100 ms. The ion mobility range was set from 0.85 to 1.30 Vs cm⁻². diaPASEF acquisition was performed with a cycle time of 0.95 s, comprising 21 mass windows per cycle with a window width of 25 Da. Collision energy was ramped as a function of ion mobility from 20 eV (1/K0 = 0.6 Vs cm⁻²) to 59 eV (1/K0 = 1.6 Vs cm⁻²).

### Generation of an isoform-resolved spectral library

Three out of four 2D-PAGE replicates were analyzed independently to generate spectral libraries using FragPipe v22 (Kong et al. 2017). For each gel we processed .d DIA files with the default diaTracer (K. Li et al. 2025) workflow to generate pseudo-MS/MS spectra. These were then searched against the human UniProt reviewed proteome with isoforms (release 2025_07_01) using the default Fragpipe workflow, with spectral-library generation and sub-mzML output enabled. This generated a fully tryptic library. The resulting sub-mzML files were subsequently searched using the FragPipe semi-tryptic second-pass workflow to generate a complementary semi-tryptic library on the same fasta file. MSBooster was disabled for the semi-tryptic search. The fully tryptic and semi-tryptic libraries were combined into one UniProt-derived library per 2D-PAGE experiment. The same procedure was repeated using a PeptideAtlas-derived FASTA (plasma, downloaded 2023-06), generating a second experimental library. We joined all per-gel spectral libraries with a predicted spectral library for human plasma proteomics that was generated using experimentally observed peptide sequences from PeptideAtlas as input. Peptide sequences were downloaded from the PeptideAtlas Human Plasma build (build 559, accessed 16 December 2023) via the SBEAMS interface, including fully tryptic, missed cleavage-containing, and semi-tryptic peptides, yielding 267,383 unique sequences. These were mapped to proteins by exact subsequence search against the UniProt human reference proteome (UP000005640, canonical sequences, Swiss-Prot only, accessed 27 March 2023) to generate a peptide-centric FASTA. *In silico* fragment ion spectra were predicted for all entries using DIA-NN (version 1.8.1), with carbamidomethylation of cysteine as a fixed modification and oxidation of methionine as a variable modification. Before concatenating the spectral libraries across gels and fasta files decoys were removed and annotation of all mapped proteins corrected to the same set of proteins per protein ID. Libraries were concatenated, for every peptide the union set of proteins and genes across libraries annotated as well as proteotypicity re-annotated based on the corrected set of genes and finally duplicated fragments removed. The resulting library columns were renamed and reformatted for quantification with DIA-NN, missing information such as termini-annotation and protein names assigned from the input fasta files. Non-human proteins added through the diaTracer workflow were also removed before running DIA-NN for quantification. Retention times, ion mobility and spectra were predicted using DIA-NN 2.3.1.

### Peptide quantification and pre-processing

This library was used with DIA-NN 2.3.1 for peptide precursor quantification, without matching between runs, unrelated runs enabled, normalisation switched off and the options --no-lib-filter and --duplicate-proteins enabled to keep the generated library as created. The outcome data was filtered by 1% FDR and peptide precursors summarised to peptides. Next, the data was scaled per peptide such that the maximum abundance of a peptide over the gel equals one. To retrieve a dataset with similar sample quality, outlier samples were detected by evaluating the identification and intensity distributions of a spiked-in internal peptide standard (OSPP) (Wang et al. 2025) and excluding samples dominated by potential contaminants, as defined peptides from the cRAP databases containing commonly used commercial reagents in laboratories, MaxQuant contaminants, the Global Proteome Machine common Repository of Adventitious Proteins (GPM-cRAP) and Cambridge Centre for Proteomics cRAP (CCP-cRAP, downloaded 06.05.2026 7:06 CET). We extended this list of potential contaminant proteins with keratins and peptides identified across at least half of any 2D-PAGE replicate. Samples with a total peptide abundance above a replicate-specific intensity distribution outlier threshold and where at least 8 of the 10 most abundant peptides originated from the potential contaminant list were excluded from further analysis. Since some of the most common plasma proteins (Albumin, complement factors) were part of these contaminant databases, we decided to flag potential contaminant proteins in our proteoform map rather than excluding them. Summing up all peptide abundances per sample (voxel in a gel) resulted in digital representations of the gels which were subdivided by up to 60,000 peptide gel representations each. For every peptide in every 2D-PAGE replicate (=spatial map) the noise level was defined as the median peptide intensity over the gel + two times the standard deviation of the noise (Immerkær 1996). Missing values were imputed using spatial information as the mean of the surrounding 8 voxels in a gel.

### Proteoform identification

Peaks per peptide were identified as local maxima with the peak_local_max algorithm from the python scikit-image package (van der Walt et al. 2014) above the noise level and at least 10% higher than any neighbouring sample (=voxel in the spatial map). Identified peaks of every peptide in a protein group (reduced to canonical UniProt Ids) were overlayed and thus proteoforms per protein group defined. Peptides which may originate from several proteins were re-annotated to a smaller list of candidate proteins if within two voxels around their identified peaks no other peptide of a candidate protein was identified. Finally, proteoforms per protein group were matched across the four replicate gels by creating graphs for every protein group with proteoforms as nodes and edges between proteoforms of different replicate gels if their distance was within a pairwise threshold (allowed voxel shift). The threshold between any two replicate gels was identified by manually measuring the shifts of all major visible spots in the gel scans. The maximum matching for a protein group was solved using node-assignment integer linear programming, decomposed into connected components solved independently with the CP-SAT solver from the python OR-tools package (Google 2026). If several possible matchings of equal size were identified, the matching with highest harmonic mean across peak peptide abundance correlations was selected as the best match. If no match could be identified in linear time, a feasible match was selected. The number of replicate gels a proteoform was identified in served as a confidence score for the proteoforms. Additionally, every proteoform landscape of a protein group was condensed into one dimension, such that it could also be matched to dSEC and IEF profiles.

## Acknowledgements

We thank our technical lab assistants Christiane Kilian and Daniela Ludwig for helping with the experiments as well as our computational team Oliver Lemke, Clemens Dierks, Dominik Bierbaum, Luise Nagel, Roza Mizrak, Vadim Farztdinov, and Ziyue Wang and Jan Gieseler for valuable discussions about the entire project.

## Author contributions

B.M.Z, T.S. and M.R planned the experiments, T.S., A.N., F.A., K.T.-T. prepared samples for 2D-PAGE, IEF and dSEC and ran preliminary experiments. K.L. ran the 2D-PAGE, T.S. performed the IEF and dSEC experiments and processed samples for LC-MS/MS. LC-MS/MS analysis was conducted by A.N. for 2D-PAGE experiments, F.A. for dSEC- and IEF-experiments and K.T.-T. during preliminary experiments. All experimental steps were supervised by T.S. Data analysis was planned and performed by B.M.Z.. T.W.S ran FragPipe analysis for spectral library generation and A.Y. designed proteoform-specificity scoring. C.G. handled data transfer, software setup and server maintenance as well as data storage. Figures for the manuscript were created by B.M.Z, T.W.S. and A.Y. The draft manuscript was written by B.M.Z, T.S., M.R. T.W.S. and L.R.S.. M.V., M.M. and M.R. supervised the study and secured funding.

